# ExoChew: An exonuclease technique to generate single-stranded DNA libraries

**DOI:** 10.1101/2023.10.02.560524

**Authors:** Krishna Patel, Chirag Lodha, Christopher Smith, Levi Diggins, Venkata Kolluru, Daniel Ross, Christopher Syed, Olivia Lewis, Rachel Daley, Bidyut K Mohanty

## Abstract

Although DNA in the genome is double-stranded, single-stranded DNA is generated during various processes including DNA replication and repair. Some single-stranded DNAs can form noncanonical structures. Various proteins bind to the single-stranded DNAs site-specifically and/or structure-specifically to regulate various DNA transactions. Because of the transient nature of single-stranded DNAs in the cell, current *in vivo* techniques may not reveal all such sequences, structures, and protein-DNA complexes. To explore such sequences and structures genome-wide, it is necessary to generate single-stranded DNA libraries. Current *in vitro* methods involve heat denaturation of libraries of double-stranded DNA fragments followed by cooling to prevent reannealing; however, a significant amount of DNA can reanneal to regenerate double-stranded DNAs. In ExoChew method, double-stranded DNA fragment libraries are enzymatically converted to single-stranded DNA libraries. Genomic DNA is sonicated to generate pools of double-stranded DNA fragments of required size. Each pool of double-stranded DNA fragments is then treated with either T7 exonuclease or *E. coli* exonuclease III. which recognize and cleave double-stranded DNA from 5’ ends or 3’ ends, generating single-stranded DNA pools, respectively. The enzymatically generated single-stranded DNA pools can be used for genome-wide studies of protein-DNA interactions and structural studies of DNA.

## Introduction

Chromosomal DNA in cellular organisms is double stranded (dsDNA); however, single-stranded DNA (ssDNA) can be formed transiently during DNA replication, transcription, telomere synthesis, recombination, and repair^1^. In addition, ssDNA is present at chromosomal ends (telomeres) where it forms a telomere loop (t-loop)^2,3^. Certain DNA sequences can generate noncanonical DNA secondary structures including DNA hairpins, cruciform, triple helix, G-quadruplexes (G4s) and intercalating motifs (i-motifs)^4,5^. These noncanonical structures are generated from non-Watson-Crick base-pairing between or among intra-strand nucleotides^4,6^.

A wide array of proteins exhibit binding affinity for dsDNA; notably a recent study identified 2604 proteins that bind dsDNA of the human genome^7^. In addition, there are various proteins that can bind to ssDNA^8^ and noncanonical DNA structures distributed throughout the genome^9^. The dsDNA- and ssDNA-binding proteins can bind DNA either site-specifically or nonspecifically^10^. Various *in vivo* analytical techniques, including chromatin immunoprecipitation (ChIP), have been used to identify genome-wide and individual protein-DNA interactions^11–13^. Typically, ChIP involves cross-linking of chromatin-bound proteins in the cell with formaldehyde, sonication, or nuclease treatment to obtain small DNA fragments bound to proteins, followed by immunoprecipitation using antibodies to specific proteins of interest^12^. This cross-linking primarily captures interactions involving dsDNA regions with very few ssDNA regions^12,13^. The number of ssDNA sequences bound to proteins *in vivo* can be very limited due to the fact that ssDNAs are generated transiently and not all such DNAs are available for protein binding at any given time in a cell or specific tissue^1^.

There are several ssDNA binding proteins in the cell^1^. Replication protein A (RPA) of mammalian cells and ssDNA binding protein (SSB) of *E. coli* bind ssDNA nonspecifically^14,15^. Several proteins are known to bind ssDNAs site-specifically or to various secondary structures generated from ssDNAs^1^. For example, hnRNP LL binds to i-motif generated from polycytosine-rich ssDNA region proximal to BCL2 oncogene promoter^16,17^. Several other hnRNPs such as hnRNP E1, hnRNP K, hnRNP A1, and hnRNP A2/B1 also bind to ssDNA sequences, specifically^18–21^. Importantly, genome-wide studies have not been conducted with most of these proteins.

To explore site-specific protein-DNA interactions genome-wide *in vitro*, ssDNA libraries must be generated from genomic DNA (gDNA). Typically, for studies involving generation of ssDNA libraries, first a dsDNA library is generated from gDNA and the library is heat denatured followed by cooling to prevent reannealing of complementary strands^22^. Although this method works well, it is possible that during the cooling process, a significant fraction of ssDNA population can undergo reannealing between complementary strands. An alternative method is to treat the dsDNA library with alkali or DMSO that denatures the dsDNA followed by neutralization of the reaction mixture^23,24^. This process can also lead to reannealing between complementary strands of DNA^24^.

We present a simple, rapid enzymatic method for the generation of ssDNA libraries from dsDNA libraries of different sizes. In this method, libraries of dsDNA fragments are treated with either T7 exonuclease^25,26^ or *E. coli* exonuclease III^27^, resulting in the generation of ssDNA libraries^28^. These exonucleases exhibit specificity for binding to dsDNA, allowing them to dissociate from the DNA strand before its hydrolysis is completed, generating ssDNA^25–28^. The T7 exonuclease enzyme hydrolyzes dsDNA from 5’ to 3’ ends from each strand of the dsDNA^25,26^. Upon reaching the midpoint of the strand, the enzyme halts, and the resultant product is an ssDNA on each side^25,26^. In contrast, *E. coli* exonuclease III works from the 3’ to 5’ ends in a progressive manner^27^. After about 30 to 45% of the dsDNA is hydrolyzed, the enzyme stops^27^. Mechanisms of action of both enzymes are illustrated in **Fig. 1A** and the use of the technique is discussed in detail in the Discussion section (**Fig. 1B**). The technique also holds potential for its application in genome-wide studies involving gDNA from single-cells. Similarly, ExoChew can also find its use in generating ssDNA from a known dsDNA fragment for a specific purpose.

**Figure 1.**
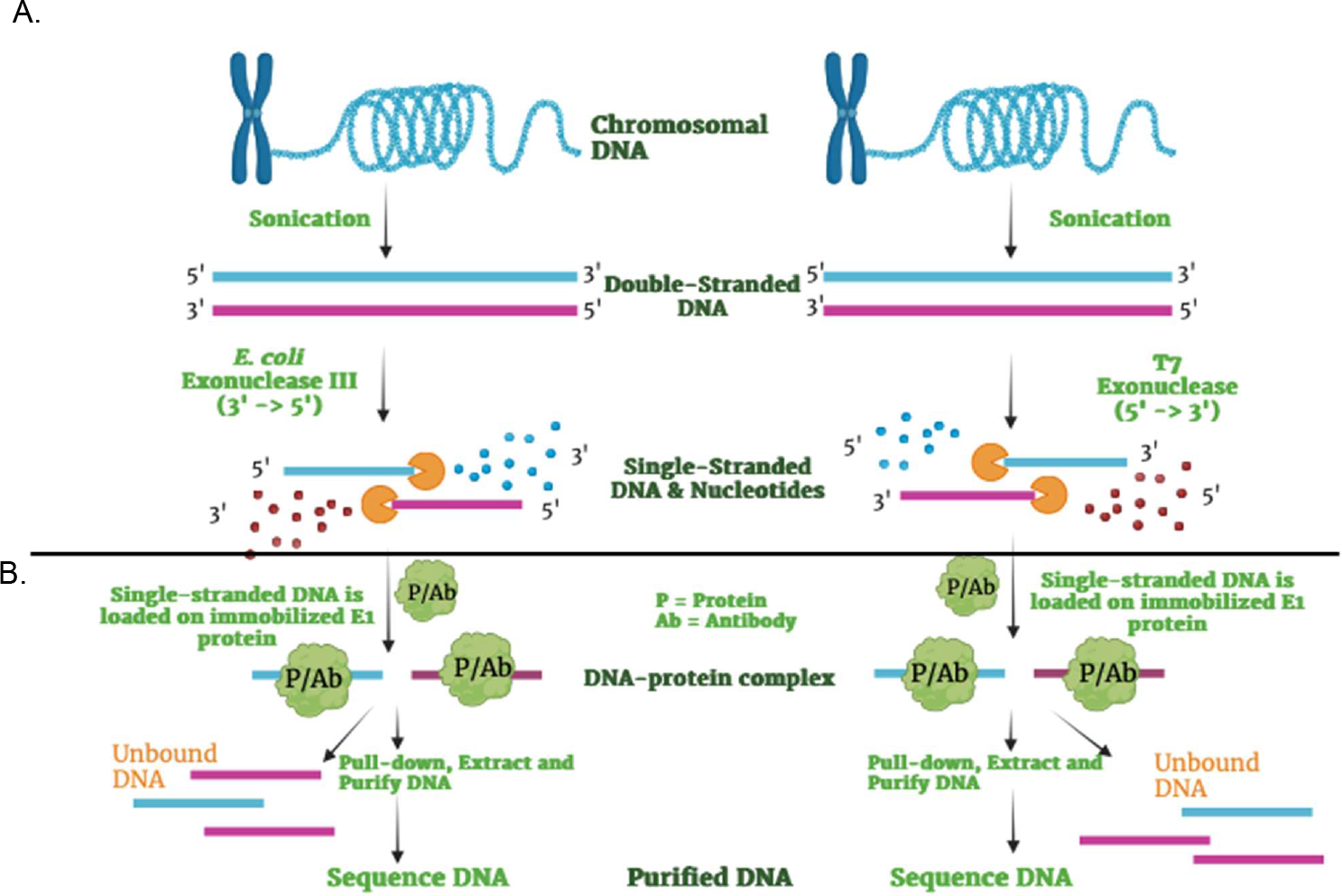
The ExoChew technique. (**A**) An illustration showing how T7 exonuclease and *E. coli* exonuclease III degrade the double stranded gDNA in either 5’ → 3’, or 3’ → 5’ directions, respectively, to form ssDNA. (**B**) The illustration demonstrates the binding of a protein or an antibody to an appropriate ssDNA sequence or its secondary structure. The DNA-protein complexes are pulled down with appropriate antibody/ligand-tagged agarose beads followed by extraction and purification of the DNA. The DNA pulled down by a protein or an antibody is then analyzed or sequenced.

## Results

### ExoChew technique

Traditionally, heat denaturation of dsDNA followed by slow cooling to room temperature or snap chilling generates ssDNA. However, reannealing of complementary DNA strands can occur with time. To circumvent this problem, we have developed ExoChew (<u>Exo</u>nuclease <u>Chew</u>ing). In this technique, as shown in **Fig. 1A**, gDNA is sonicated (using an ultrasonicator or a manual sonicator) to generate a pool of dsDNA fragments of appropriate size. The pool is divided into three sub-pools, with the first sub-pool treated with T7 exonuclease, the second with *E. coli* exonuclease III, and the third used as a control. T7 exonuclease recognizes dsDNA and catalyzes the removal of mononucleotides in the 5’ -> 3’ direction. In contrast, *E. coli* exonuclease III recognizes dsDNA and removes nucleotides in the 3’ --> 5’ direction. Each enzyme will generate a pool of ssDNAs, which can then be screened for identifying ssDNA sequences that bind to specific proteins or for generating specific structures (e.g., G4s and iMs) followed by their interaction with specific proteins or antibodies.

### ExoChew of 100-base pair DNA ladder

To determine the efficacy of the exonucleases, we first treated a commercially available 100 base pair (bp) DNA ladder with each of the two enzymes. As shown in **Fig. 2A**, the 100 bp ladder was treated with T7 exonuclease and analyzed by non-denaturing polyacrylamide gel electrophoresis (PAGE). To determine if the T7 exonuclease generated single-stranded DNA from the dsDNA ladder, we incubated the untreated and T7 exonuclease-treated DNA samples with *E. coli* single-stranded DNA binding protein (SSB) and analyzed the samples by PAGE. As shown in **Fig. 2A**, treatment of the 100bp dsDNA ladder with T7 exonuclease generated a completely different pattern of DNA fragments from the untreated DNA. Further, whereas incubation of untreated 100-bp ladder with SSB did not result in any significant change in the electrophoretic pattern of the DNA bands, incubation of the T7 exonuclease-treated DNA ladder with SSB caused slow mobility of the DNA fragments. This suggests that T7 exonuclease treatment generated ssDNA fragments of the 100-bp DNA ladder.

**Figure 2.**
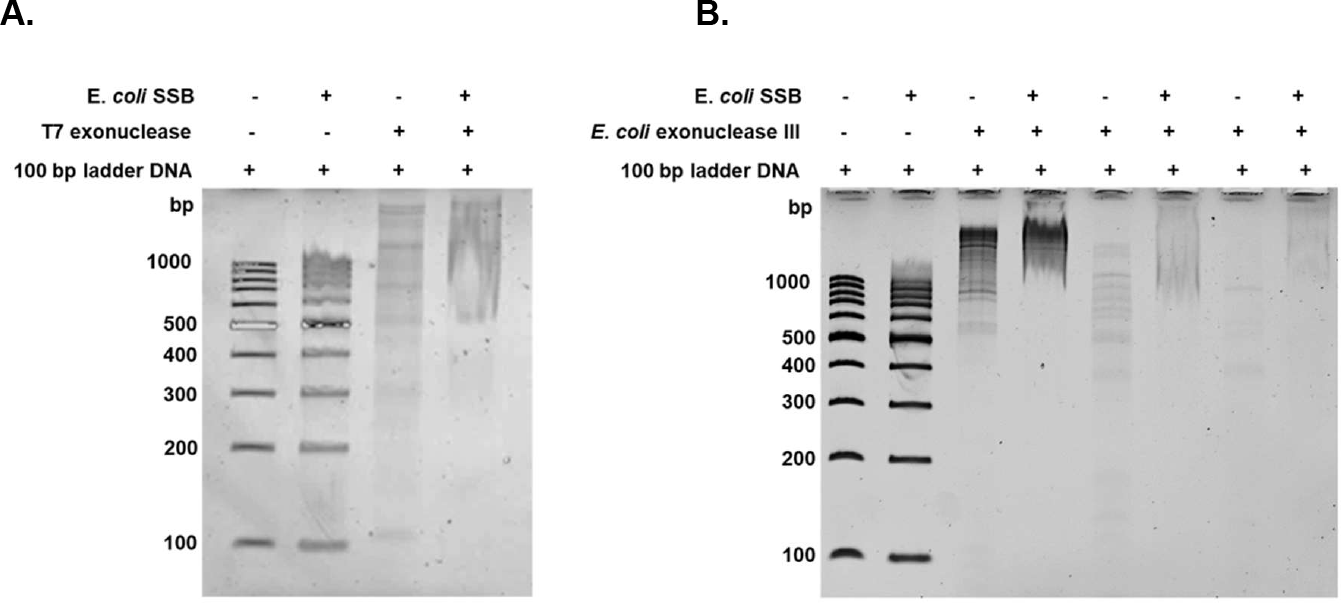
Treatment of 100 bp ladder with T7 exonuclease (A) and *E. coli* exonuclease III (B) and binding by single-stranded binding protein (SSB). (**A**) A 6% polyacrylamide gel showing the difference in SSB binding between 100 bp DNA ladder without T7 exonuclease treatment and upon treatment with T7 exonuclease. Addition of SSB to the Exochew (by T7 exonuclease) product of the DNA ladder shows slower migration of DNA in the gel. Experimental conditions are described in the Materials and Methods section. (**B**) A 6% polyacrylamide gel showing the difference in SSB binding between 100 bp DNA ladder without *E. coli* exonuclease III treatment and upon *E. coli* exonuclease III treatment. Experimental conditions are described in the Materials and Methods section.

We also treated the 100-bp dsDNA ladder with *E. coli* exonuclease III enzyme. As shown in **Fig. 2B**, treatment of the 100-bp ladder with the nuclease changed the pattern of DNA fragments in the gel and the changes were much more discernible with time. The untreated and *E. coli* exonuclease III-treated DNA samples were then incubated with SSB. Whereas SSB treatment did not affect the mobility of untreated 100-bp DNA fragments in a nondenaturing PAGE gel, SSB treatment of the nuclease-treated DNA samples showed significant slowdown of their migration in the gel. All these results strongly suggest that T7 exonuclease and *E. coli* exonuclease III efficiently generated ssDNA fragments of the 100-bp DNA ladder.

### ExoChew of human genomic DNA

As with 100-bp DNA ladder, human genomic DNA from A549 cells was tested to determine the success of ExoChew technique using T7 exonuclease. As shown in **Fig. 3A**, the human gDNA derived from A549 cells was fragmented using the ultrasonicator. The dsDNA fragment pool was then treated with T7 exonuclease (**Fig. 3B**). Untreated and T7-exonuclease-treated dsDNA fragments were incubated with SSB followed by fractionation in a PAGE gel. As shown in **Fig. 3C**, whereas SSB did not show any change in the mobility of dsDNA pool upon gel fractionation, it changed the mobility of the T7-exonuclease-treated DNA pool. The results show that T7 exonuclease successfully converted the pool of A549 human dsDNA fragments into ssDNAs.

**Figure 3.**
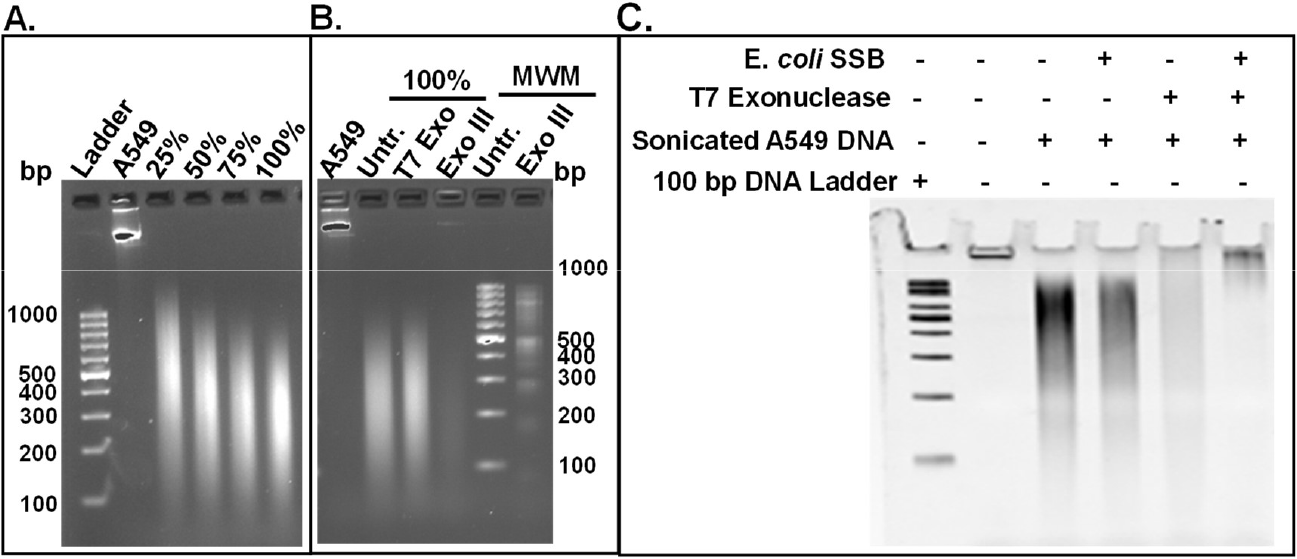
ExoChew and SSB binding with A549 human gDNA. (**A**) Agarose gel showing the sonication products of gDNA from A549 cells to different stages of completion. 100% shows the completed sonication products of the dsDNA. (**B**) Polyacrylamide gel electrophoresis showing ExoChew products of A549 gDNA fragment pools. Lanes are: A549 gDNA, sonicated A549 gDNA (untr.), T7 exonuclease-treated dsDNA pool (T7 Exo), *E. coli* Exonuclease III-treated DNA pool (Exo III), and 100-bp DNA ladder (MWM, Untr. and Exo III). (**C**) PAGE gel showing binding of *E. coli* SSB to ExoChew products of A549 DNA. T7 exonuclease treated DNA pools incubated with SSB moved slowly in a polyacrylamide gel, in contrast to the dsDNA pool that was not treated with exonuclease. In contrast, dsDNA pool did not show any change in its mobility in the presence of SSB.

### ExoChew of mouse genomic DNA

To determine the efficacy of ssDNA generation in the dsDNA pool from mouse gDNA, we sonicated mouse gDNA derived from NMuMG cells using ultrasonicator to either 200-bp fragments or 350-bp fragments. As shown in **Fig. 4A**, smears of DNA were generated upon fragmentation of the gDNA. The dsDNA pools were treated with T7 exonuclease or *E. coli* exonuclease III as described in Materials and Methods. The untreated, sonicated dsDNA fragments and exonuclease treated DNA samples are shown in **Fig. 4B**. The samples were then tested for their binding to SSB. As shown in **Fig. 4C**, the pattern of the untreated dsDNA smear did not change much upon addition of SSB. However, the T7 exonuclease-treated DNA fragments and *E. coli* exonuclease III-treated DNA fragments showed slowdown in their migration in the gel in the presence of SSB (**Fig. 4C**). These results strongly suggest that T7 exonuclease and *E. coli* exonuclease III enzymes generated ssDNA pools from the dsDNA pools of mouse gDNA.

**Figure 4.**
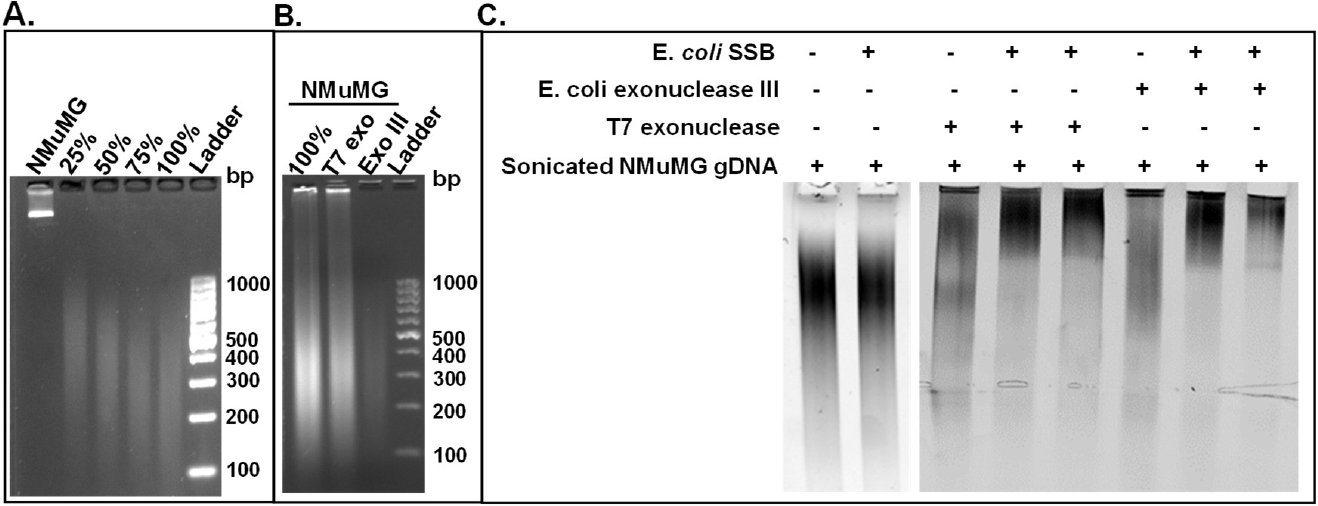
ExoChew and SSB binding with NMuMG mouse gDNA. (**A**) Agarose gel showing the sonication products of NMuMG (mouse) DNA. 100% shows the completion of sonication of the dsDNA. (**B**) Agarose gel showing ExoChew products of sonicated NMuMG gDNA. Lanes: sonicated NMuMG dsDNA, T7 Exo product, and Exo III product. (**C**) Polyacrylamide Gel Electrophoresis showing SSB binding to ExoChew products of NMuMG DNA. Lanes (from left to right): sonicated NMuMG dsDNA (100%), sonicated NMuMG dsDNA + SSB, T7 Exonuclease-treated NMuMG DNA, T7 Exonuclease-treated NMuMG DNA with two different concentrations of SSB, *E. coli* Exonuclease III-treated NMuMG DNA, and *E. coli* Exonuclease III-treated NMuMG DNA with two different concentrations of SSB. Whereas dsDNA pools did not show any change in their mobility in the absence or presence of SSB, dsDNA pools treated with T7 Exonuclease and *E. coli* Exonuclease III moved slowly in comparison to similar pools without SSB treatment.

## Discussion

Single-stranded DNA libraries generated by ExoChew method can be used to conduct genome-wide search for ssDNA-protein interactions. The ssDNA libraries can also be modified appropriately to generate libraries of specific DNA secondary structures in the total pool. For example, by modifying the pH of the buffer containing the ssDNA pool, libraries of i-motifs or G-quadruplex secondary structures^22,29^ can be generated. We are currently working on generating such specific pools of DNA for genome-wide studies.

Two potential problems may be encountered in ExoChew technique: (1) The length of the DNA is reduced by half at the end of the conversion of dsDNA to ssDNA by the exonucleases. We address the issue by conducting ExoChew with two exonucleases in two separate pools of dsDNA (**Fig. 1A**). Whereas the T7 exonuclease cleaves dsDNA in 5’ → 3’ direction, the *E. coli* exonuclease III cleaves dsDNA in 3’ → 5’ direction. After exonuclease treatment, the two pools are not mixed; they are treated in the same manner until the end (**Fig. 1B**). (2) Since there are multiple copies of each chromosome and sonication generates fragments of these chromosomes randomly, the dsDNA fragments and, thus, the ssDNA fragments may have some small overlaps that can cause annealing between complementary sequences. This problem is addressed in two ways: (i) since the exonuclease is still present and active in the reaction mix, it will cleave the dsDNAs generated by partial annealing between two ssDNAs generating all ssDNA molecules; (ii) when exonuclease is added to dsDNA library for ExoChew reaction, a specific protein that binds to ssDNAs or a ligand or an antibody that binds to a specific DNA structure can be added simultaneously so that the protein-DNA complexes, the ligand-DNA complexes, or antibody-DNA complexes are formed in real time as the ssDNAs are being generated by ExoChew.

We do not know yet if we will lose some sequences in ExoChew technique (e.g., telomeres or other DNAs containing multiple repeats of short sequences). We will examine this in two ways: (i) A dsDNA pool and its ExoChew products will be sequenced. A comparison of the two libraries will answer this question. (ii) As mentioned in a previous section, we will mix a protein, a ligand, or an antibody of choice with a dsDNA library along with an exonuclease (T7 exonuclease or *E. coli* exonuclease III) at the same time to simultaneously initiate ExoChew and generate iM-protein/antibody complexes. The bound DNAs will be sequenced. In parallel, we will sequence the protein/antibody-bound ssDNAs from ssDNA pool generated by traditional heat denaturation method.

In summary, ExoChew is a simple enzymatic technique to generate libraries of ssDNA fragments from a library of dsDNA fragments.

## Materials and Methods

### Cell lines, plasmids, antibodies

Mouse cell line NMuMG and human cell line A549 cells (obtained from Dr. Philip H. Howe, Medical University of South Carolinas, Charleston, SC) were grown in DMEM with a high glucose medium containing 10% FBS (Gibco).

### Chromosomal DNA preparation

Mammalian cells were grown in appropriate medium (previous section) and harvested by trypsinization followed by washing with PBS or by adding 10 mM Tris (pH 7.4), 1 mM EDTA (pH 8), and 1% SDS. Chromosomal DNA was prepared using a Quick-DNA Midiprep Plus Kit (Zymogen; D4075). DNA amount was measured using a NanoVue spectrophotometer (GE NanoVue Plus).

### Generation of double-stranded DNA library from genomic DNA

#### Sonication in Covaris

5-15 µg of gDNA in 500 µL of TE buffer was sonicated in a Covaris M220 Focused-ultrasonicator to either 200 bp or 350 bp size according to the manufacturer’s instructions. The ultrasonication parameters were set as follows: Power [75.0], Duty Cycle [20.0], Cycles per Burst [200], and Time [1 minute]. Samples were collected at 25, 50, 75, and 100% sonication. The fragmented DNA samples were analyzed in a 1.5% agarose gel in TBE buffer, stained with red dye, and visualized in Nucleic Acid gels channel using an iBright 1500 Imaging System (ThermoFisher).

#### Manual sonication

For manual sonication, the DNA samples were fragmented using a QSonic sonicator at a power of 70% with 30 second pulses and with 1 minute intervals on ice. The fragmented DNA samples were analyzed in a 1.5% agarose gel in TBE buffer and visualized using an Invitrogen iBright 1500 Imaging System (ThermoFisher).

### ExoChew method to generate single-stranded DNA pools from dsDNA pools

Fragmented dsDNA pools were treated with either T7 Exonuclease or *E. coli* Exonuclease III to generate ssDNA pools according to manufacturer’s instructions (New England Biolabs). Briefly, 1 µg dsDNA pool was treated with 10 units of T7 exonuclease (M0263) in 50 µL volume. The reaction was carried out at 25°C for 30 min and stopped by 11 mM EDTA. For *E. coli* Exo III reaction, 5 µg dsDNA pool was treated with 50 units of *E. coli* exonuclease III (M0206) in 50 µL volume. The reaction was carried out at 37°C for 30 min and stopped by 11 mM EDTA. The reaction products were first analyzed by 2% agarose gel electrophoresis. To confirm that dsDNA was converted to ssDNA, 200-300 ng of dsDNA and its exonuclease products were tested for their binding to *E. coli* SSB as described below. T7 exonuclease and *E. coli* exonuclease III reactions were scaled up depending on the requirements of ssDNA.

### Single-Stranded DNA Binding Protein (SSB) binding to ssDNA

Conversion of dsDNAs to ssDNAs was determined by their binding to SSB (MCLAB - Molecular Cloning Laboratories, CA). DNA samples were mixed with 1, 2, and 3 µL of SSB (5 µg/µL), incubated at 37°C for 5 min, and fractionated in a 6% polyacrylamide gel at 80 volts. The gels were visualized by Invitrogen iBright1500 Imaging System (ThermoFisher).

## Acknowledgements

The authors thank VCOM-Carolinas for providing financial support for the entire project.

